# Relationships of hematocrit concentration with dementia from a multiethnic population-based study

**DOI:** 10.1101/2024.08.15.608190

**Authors:** David J. Roh, Minghua Liu, Kevin Strobino, Stephanie Assuras, Vanessa Guzman, Bonnie Levin, Steven Spitalnik, Tatjana Rundek, Clinton B. Wright, Mitchell S. V. Elkind, Jose Gutierrez

**Author notes:** <u>Correspondence:</u> David Roh, MD, 177 Fort Washington Ave, 8-GS300, New York, NY 10032, Jose Gutierrez, MD/MPH, 710 W 168^th^ Street, suite 639, New York, NY 10032. David Roh, MD and Jose Gutierrez, MD/MPH take full responsibility for the data, the analyses and interpretation. These authors have full access to all of the data and have the right to publish any and all data separate and apart from any sponsor. The manuscript has been read and approved by all authors. This work has not been submitted or reported previously and represents the authors’ original work. All authors have agreed to conditions noted on the authorship agreement form. This study was approved by the IRB as stated in the Methods section.

## Abstract

**Objective:** Red blood cell (RBC) concentration impacts cerebrovascular disease, yet it is unclear whether RBC concentrations relate to dementia risk, particularly in racially/ethnically diverse cohorts. We investigated whether RBC concentrations associate with incident dementia risk in a diverse population of stroke-free individuals and explored whether cerebral small vessel disease (CSVD) mediates this relationship.

**Methods:** A longitudinal observational analysis was performed using a population-based cohort of stroke-free, older adult participants (>50 years) from the Northern Manhattan Study (NOMAS) enrolled between 2003-2008. Participants received baseline hematocrit testing, MRI neuroimaging, and cognitive assessments at baseline and long-term follow-up. Associations of baseline hematocrit as a categorical variable (low, normal [reference], and high based on laboratory reference levels) with incident dementia were assessed using Cox models adjusting for relevant covariates. Separate analyses investigated whether MRI CSVD mediated these relationships.

**Results:** We studied 1207 NOMAS participants (mean age 71±9 years, 60% female, 66% Hispanic). Mean hematocrit was 41.2% (±3.8) with 16% of participants developing incident dementia. Lower hematocrit associated with increased dementia risk (adjusted hazard ratio 1.81 [1.01-3.23]) after adjusting for age, sex, race/ethnicity, education, APOE status, and comorbidities. High hematocrit was not associated with dementia risk. No interactions by sex or race/ethnicity were seen and baseline CSVD did not mediate relationships between hematocrit and dementia.

**Conclusions:** Low hematocrit associated with dementia risk in our diverse population cohort. Further work is needed to assess mechanisms behind anemia’s relationship with dementia to assess whether this can serve as a trackable, preventable/treatable risk factor for dementia.

## INTRODUCTION

Red blood cell (RBC) concentrations are known to impact cerebrovascular disease incidence and outcomes. This relationship appears to also apply to dementia as low RBC concentrations/anemia, a prevalent condition in the elderly^1^, has been identified as an independent risk factor for incident dementia^2^. However, separate studies have identified that these relationships exist across a range of RBC concentrations, specifically with both low and high RBC concentration extremes associating with dementia risk^3,4^. It is unclear whether these findings are generalizable to multi-ethnic communities who have different risks for both anemia and polycythemia. Furthermore, underlying mechanisms for these relationships are unknown. It is currently posited that both low and high RBC concentrations could directly play a role through hypoxic/ischemic and microthrombotic cerebral insults, respectively^5–7^. We and others have separately identified that both low and high RBC concentrations associate with asymptomatic, covert cerebral small vessel disease across a variety of disease conditions^8–15^, creating a premise that ischemia or microthrombosis may indeed play a role in dementia risk. While it is known that neuroimaging evidence of covert cerebrovascular disease increases the risk of dementia^16–19^, it remains to be determined whether this mediates associations of RBC concentration and dementia. Thus, we sought to investigate the hypothesis that RBC concentration is related to incident dementia risk and that these relationships would be mediated by covert cerebral small vessel disease in a multi-ethnic, stroke-free, population-based cohort study.

## METHODS

### Northern Manhattan Study (NOMAS)

We included NOMAS participants enrolled in an MRI sub-study between 2003-2008. Participants enrolled were stroke-free individuals over 50 years old from the Northern Manhattan community that were either NOMAS participants or unrelated household members^20,21^. Participants with available baseline RBC laboratory assessments, MRI imaging, and follow-up cognitive assessments were assessed. Participants with dementia at baseline were excluded from analyses (figure 1).

**Figure 1:**
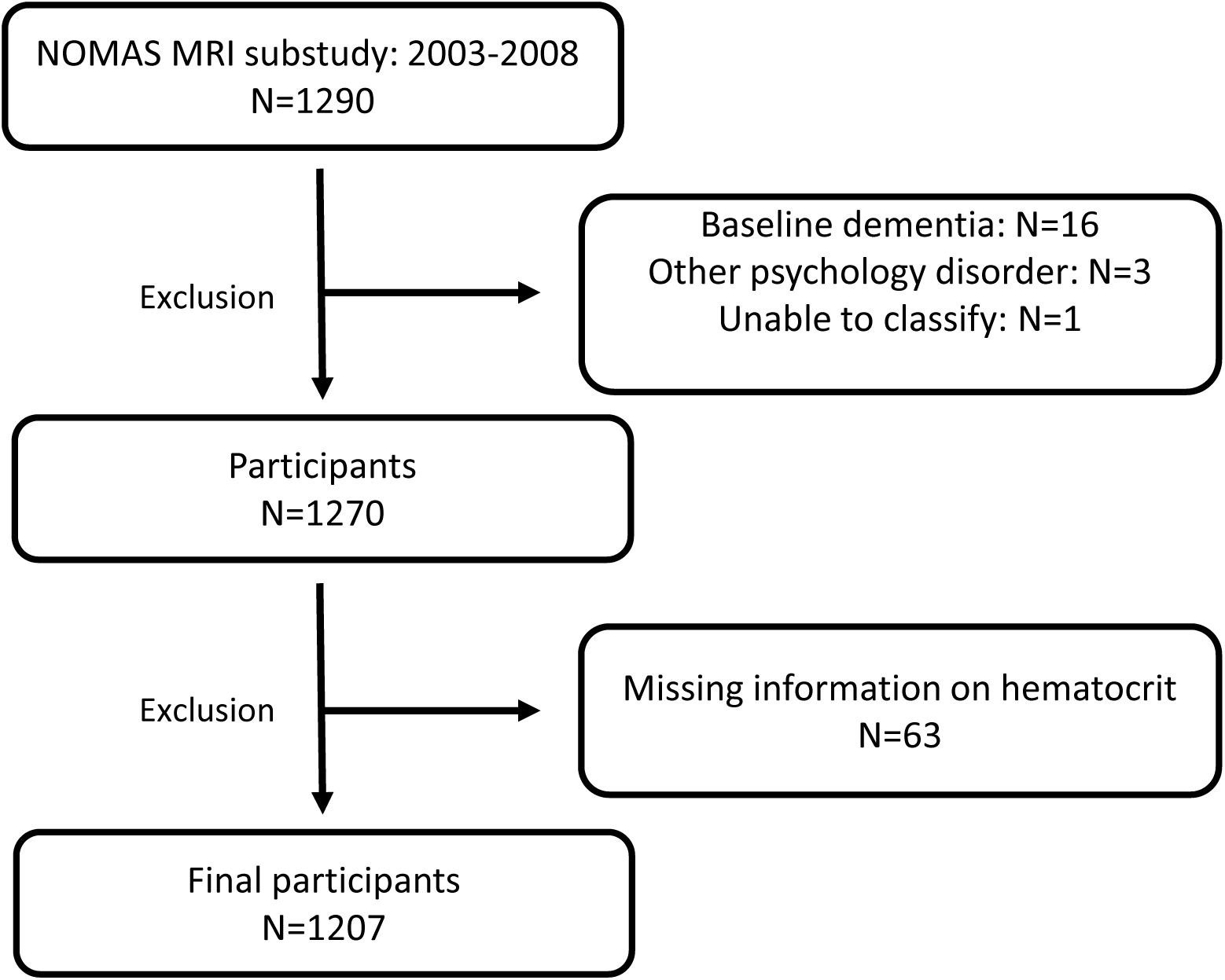
Patient inclusion and exclusion. Legend: NOMAS: Northern Manhattan Study, MRI: magnetic resonance imaging

### RBC concentration assessment

Hematocrit at the baseline visit was used as the assessment of RBC concentration. Given potential non-linear relationships of hematocrit with outcomes of interest, hematocrit was primarily assessed as a categorical variable based on reference ranges of the NOMAS core laboratory (normal vs low vs high [male normal range 37-48%; female normal range 34-43%]). Hematocrit was secondarily assessed as a continuous exposure variable. Separate analyses were performed using hemoglobin as a surrogate assessment of RBC concentration. Hemoglobin was assessed as a categorical variable similarly based on laboratory reference ranges (male normal range: 12.6-17g/dL; female normal range: 12-15.8g/dL).

### Cognitive assessments and outcome definition

Participants underwent structured questionnaires and neuropsychological assessments by trained research staff in either English or Spanish to assess cognitive function as previously described^21^. Assessments occurred at baseline with two additional follow-up visits at roughly 5-year intervals. Incident dementia was assessed as our primary outcome. Dementia was adjudicated by a multidisciplinary team, including a neurologist and a neuropsychologist, by consensus. Diagnosis was based on available clinical data including repeated neuropsychological testing, functional assessments (Informant Questionnaire of Cognitive Decline in the Elderly, Older Americans Resources and Services IADLs, Functional Assessment Scale and/or Clinical Dementia Rating [CDR] score), annual Telephone Interview for Cognitive Status (TICS) score, annual medication history (including newly self-reported dementia medications), self-reported diagnosis of dementia or self-reported “cognitive failures,” and medical records from participants who followed at our institution for their care^21^.

### MRI neuroimaging

Baseline standardized MRIs were obtained for these NOMAS participants on a dedicated 1.5T research MRI (Philips Medical Systems). MRI acquisition was performed concurrently with baseline hematocrit laboratory assessments. A composite assessment of small vessel disease severity was utilized (severe: presence of 2-4 small vessel disease markers vs non-severe: 0-1 small vessel disease markers). Alternatively, we separately assessed presence of chronic covert lacunar ischemic infarcts, severe small perivascular space burden (SPVS, defined as upper quintile), presence of severe white matter hyperintensity burden (defined as upper quintile), and cerebral microbleeds as mediating factors of interest. These markers were assessed using previously described methods via a central neuroimaging analysis core blinded to study outcomes^20^.

### Statistical analysis

Participant characteristics were compared among low, normal, and high range hematocrit. Relationships of hematocrit, defined as a categorical variable (low, normal, high) with incident dementia were assessed in primary analyses using Cox regression models. Additional models were assessed analyzing hematocrit as a continuous variable. Given potential non-linear associations of hematocrit with dementia^3,4^, separate models for hematocrit as a continuous variable were performed using restricted cubic splines regression fitting. Fine and Gray regression was performed to calculate the cumulative incidence risk per hematocrit categorical group. In secondary analyses, we assessed whether covert small vessel disease on MRI mediated the relationship of hematocrit with incident dementia. All models were adjusted for age, sex, race/ethnicity, hypertension, dyslipidemia, diabetes, smoking status, renal function (as assessed using eGFR), education attainment, APOE status, and total cranial volume. We investigated statistical interactions by sex, race/ethnicity, and APOE status. Finally, mediation analyses were performed to estimate whether small vessel disease neuroimaging markers were the mediator for hematocrit’s relationship with dementia by regressing all 3 variables together. Statistical significance was assessed at p-value <0.05. Analyses were performed using SAS (version 9.4, SAS Institute, Cary, NC).

### Standard protocol approvals and patient consents

The NOMAS protocol was approved by the Institutional Review Boards at Columbia University Irving Medical Center and the University of Miami. Informed consent was obtained from all participants.

## RESULTS

Data from 1207 stroke-free participants from NOMAS with available hematocrit and cognitive outcomes were assessed (figure 1). Baseline characteristics are shown in table 1. The mean age of our cohort was 71, 60% were female, and 66% were Hispanic. The mean hematocrit was 41.2%. The majority of participants had hematocrits within the normal reference range (83%) with low and high hematocrit identified in 5% and 12%, respectively. We identified that participants with low hematocrit were more likely to be older, non-Hispanic Black, and diabetic compared to normal and high hematocrit participants (table 1). There were no notable intergroup differences in baseline cognitive assessments, or years of education between low, normal, and high hematocrit groups.

**Table 1:** Baseline characteristics of NOMAS participants with normal, low, and high hematocrit.

|  |  |  |  |  |  |
| --- | --- | --- | --- | --- | --- |
|  | **Overall**<br>**NOMAS cohort**<br>**N= 1207** | **Normal Hct**<br>**N=1000** | **Low Hct**<br>**N=59** | **High Hct**<br>**N=148** | **P-value** |
| **Age:** mean (SD) | 70.5 (8.9) | 70.3 (9.0) | 73.1 (8.5) | 70.4 (8.6) | 0.055 |
| **Female:** N (%) | 722 (59.8) | 570 (57.0) | 32 (54.2) | 120 (81.1) | <0.001 |
| **Race:** N (%) |  |  |  |  | 0.002 |
| Non-Hispanic White | 205 (17.0) | 172 (17.2) | 10 (16.9) | 23 (15.5) |  |
| Non-Hispanic Black | 203 (16.8) | 162 (16.2) | 21 (35.6) | 20 (13.5) |  |
| Hispanic | 799 (66.2) | 666 (66.6) | 28 (47.5) | 105 (70.9) |  |
| **Medical History:** N (%) |  |  |  |  |  |
| Hypertension | 947 (78.5) | 776 (81.9) | 50 (84.7) | 121 (81.8) | 0.250 |
| Diabetes | 309 (25.6) | 260 (26.0) | 24 (40.7) | 25 (16.9) | 0.001 |
| Dyslipidemia | 977 (80.9) | 806 (80.6) | 52 (88.1) | 119 (80.4) | 0.353 |
| **Smoking:** N (%) | 141 (11.7) | 107 (10.7) | 7 (11.97) | 27 (18.2) | 0.029 |
| **Hematocrit (%):** mean (SD) | 41.2 (3.8) | 41.1 (3.0) | 33.2 (2.2) | 45.8 (2.8) | <0.001 |
| **Cognitive assessments:** |  |  |  |  |  |
| Number of assessments: median (IQR) | 2 (2-3) | 2 (2-3) | 2 (1-2) | 2 (2-3) | 0.002 |
| Time to first follow-up (years): median (IQR) | 0.93 (0.78-1.09) | 0.93 (0.77-1.09) | 0.94 (0.8-1.06) | 0.91 (0.80-1.06) | 0.515 |
| Follow up (years): median (IQR) | 9.6 (5.1-12.5) | 10.4 (5.1-12.6) | 5.1 (4.9-10.7) | 5.7 (5.1-12.3) | 0.001 |
| Years of education: mean (SD) | 9.7 (5.2) | 9.7 (5.2) | 10.5 (5.1) | 9.1 (5.0) | 0.157 |
| Baseline cognitive domain Z-score: mean (SD) | -0.006 (0.718) | -0.001 (0.725) | 0.019 (0.643) | -0.049 (0.70) | 0.727 |
| Baseline MMSE: mean (SD) | 26.7 (3.3) | 26.8 (3.2) | 27.3 (3.2) | 26.4 (3.6) | 0.197 |
| Incident Dementia: N (%) | 188 (15.6) | 152 (15.2) | 14 (23.7) | 22 (14.9) | 0.207 |
| **Baseline MRI findings:** N (%) |  |  |  |  |  |
| Chronic covert lacunar infarct | 214 (17.7) | 175 (17.5) | 13 (22.0) | 26 (17.6) | 0.674 |
| Small perivascular space | 74 (6.1) | 64 (6.4) | 4 (6.8) | 6 (4.1) | 0.528 |
| NOMAS: Northern Manhattan Study; SD: standard deviation; Hct: hematocrit; MMSE: mini mental state evaluation |  |  |  |  |  |

Amongst our entire cohort, incident dementia was adjudicated in 16% over a median total follow-up time of 9.6 years. When assessing hematocrit as a categorical variable (normal, low, high) using laboratory reference ranges set by our centralized laboratory, we identified that patients with low hematocrit had an increased risk of incident dementia when referenced to participants with normal hematocrit concentrations (adjusted hazard ratio 1.81 [1.01-3.23], p=0.04). No associations of high hematocrit category with dementia (adjusted hazard ratio 0.70 [0.43-1.15], p=0.16) were seen. When assessing the relationship of baseline hematocrit as a continuous linear variable with incident dementia, we again identified that lower hematocrit associated with increased risk of dementia (adjusted hazard ratio 0.95 per 1% change in hematocrit [0.91-0.99]; p=0.04). We did not identify significant U-shaped relationships of hematocrit extremes with incident dementia (chi-square: 2.08; p=0.56). The Fine and Gray test identified that participants with lower hematocrits had higher probability of dementia over shorter periods of follow-up time compared to normal and high hematocrit groups, but these differences were not statistically significant (p=0.20; see figure 2). Separate models utilizing hemoglobin instead of hematocrit as the assessment of RBC concentration yielded similar associations of low, but not high, hemoglobin categories with incident dementia risk (low hemoglobin adjusted HR 1.54 [1.04-2.28], p=0.03; high hemoglobin adjusted HR 0.70 [0.17-2.87], p=0.62).

**Figure 2:**
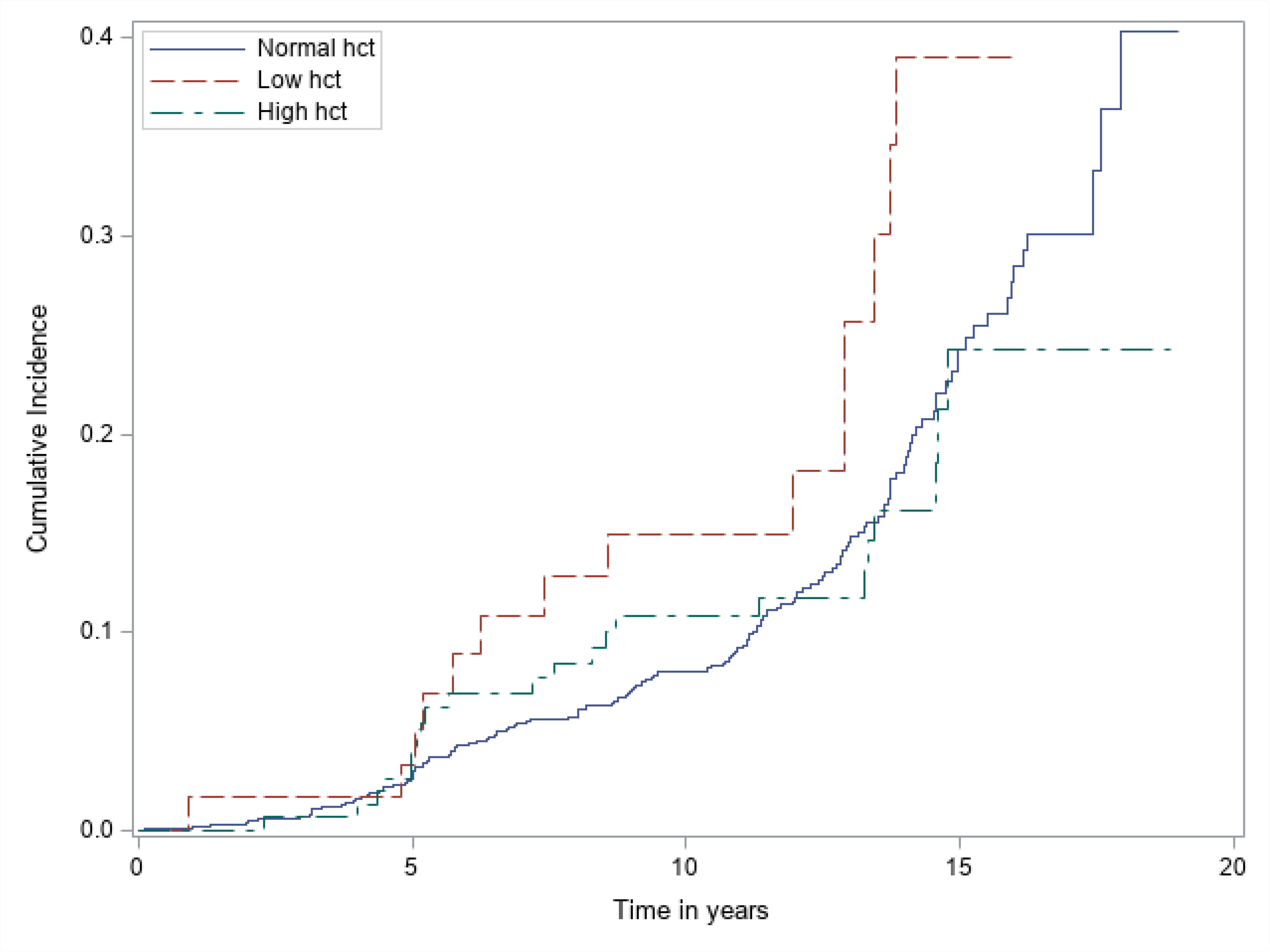
Cumulative incidence of dementia by different hematocrit groups (low vs normal vs high) Legend: Hct: hematocrit

In our MRI analyses of baseline chronic, covert cerebral small vessel disease markers, we identified severe cerebral small vessel disease burden in 12% of the overall cohort. Assessments of individual cerebral small vessel disease markers by hematocrit categories can be seen in table 2. We identified that participants with low hematocrit were more likely to have baseline MRI evidence of white matter disease burden, however differences were non-significant. In our mediation analyses, we did not identify clear evidence that either composite severe small vessel disease presence or separate, individual small vessel disease markers (chronic covert lacunar ischemic infarcts, SPVS, white matter hyperintensity, cerebral microbleeds) mediated the relationship of low hematocrit with incident dementia (supplemental figure).

**Table 2:** Prevalence of chronic covert cerebral small vessel disease MRI markers by hematocrit group, MRI cerebral small vessel disease markers in NOMAS participants with normal, low, and high hematocrit.

| **Table 2: MRI cerebral small vessel disease markers in NOMAS participants with normal, low, and high hematocrit** |  |  |  |  |  |
| --- | --- | --- | --- | --- | --- |
|  | **Overall**<br>**NOMAS cohort** | **Normal Hct** | **Low Hct** | **High Hct** | **P-value** |
| **Lacunar infarct** | 74 (6) | 64 (6) | 4 (7) | 6 (4) | 0.53 |
| **Small perivascular space** | 220 (19) | 180 (19) | 9 (16) | 31 (22) | 0.59 |
| **White matter hyperintensity** | 232 (19) | 187 (19) | 17 (29) | 28 (19) | 0.16 |
| **Cerebral microbleed** | 89 (8) | 69 (7) | 6 (10) | 14 (10) | 0.46 |
| **Severe composite small vessel disease** | 129 (12) | 103 (11) | 9 (16) | 17 (12) | 0.55 |
| NOMAS: Northern Manhattan Study; Hct: hematocrit |  |  |  |  |  |

## DISCUSSION

In our large population-based study of ethnically diverse, non-stroke participants over 50 years old with long-term follow-up, we identified independent relationships of lower hematocrit with incident dementia after adjusting for known risk factors of dementia. While we evaluated hematocrit across its entire range, it was notable that we did not identify additional relationships of higher hematocrit with dementia. And despite our prior published findings identifying relevant relationships of lower hematocrit concentrations with MRI evidence of asymptomatic, covert cerebral small vessel disease^8^, these MRI biomarkers did not appear to be primary drivers/mediators for low hematocrit’s relationship with dementia.

Previous population cohorts have reported U-shaped relationships of RBC concentrations at both extremes (ie., anemia and relative polycythemia) with incident dementia^3,4^. Yet, despite having similar dementia prevalence in our cohort compared to these prior studies, we were only able to replicate lower hematocrit’s relationship with dementia. It is worth noting that prior studies which identified U-shaped relationships of both RBC concentration extremes with dementia did see greater effect estimates for dementia risk in lower RBC concentration groups compared to higher ones^3,4^. Thus, our data may add to this literature by providing evidence that lower hematocrit plays a more pervasive, generalizable role than higher hematocrit on dementia risk. While we did not identify relevant interactions of hematocrit with race/ethnicity and adjusted for baseline medical disease characteristics, it could be posited that our diverse cohort, comprised primarily of Hispanic participants with higher prevalence of baseline cardiac and cerebrovascular risk factors is more vulnerable to the impacts of low RBC concentrations. Prior multi-ethnic cohorts of Black participants have reported similar anemia-dementia relationships without interactions of race and anemia^2^.

Current mechanisms and drivers for lower RBC concentration’s relationship to dementia remains unclear. Because RBCs are critical for cerebral oxygen delivery, it has been posited that anemia creates risk for cerebral hypoxia, inflammation, and even ischemia, all factors that may lead to neuronal injury and dementia. To this extent, we had previously identified in a cross-sectional analysis of our NOMAS cohort that lower hematocrit concentrations associate with increased risk of chronic MRI cerebrovascular disease markers: covert lacunar infarcts and SPVS. Though this supported the idea that low hematocrit’s relationship with dementia could be mediated by asymptomatic, covert neuroimaging biomarkers of cerebral small vessel disease, this was not recapitulated in our mediation analyses. This could suggest that anemia’s impact on dementia is unrelated to covert cerebral small vessel disease burden as other studies (albeit in participants of primarily European descent) did not identify that anemia relates to covert infarcts. Instead, these studies identified that anemia has overlapping relevance with cerebral perfusion (unavailable in our study), an MRI neuroimaging biomarker known to be a risk factor for dementia^3,22^. But, it should be noted that our baseline MRI acquisition was performed in conjunction with our hematocrit assessments, which preceded incident dementia diagnosis by several years. Thus, it is possible that there still may be a mediating role of covert cerebral small vessel disease on our findings that simply was not identified given our study’s inherent concurrent baseline laboratory/MRI assessment design. It is feasible that dynamic changes in covert cerebral small vessel disease burden occurs over time in anemic patients which would be better assessed with serial longitudinal MRI imaging. Because mediation analyses necessitate a temporal ordering of exposure variable, mediator, or outcome, our study’s acquisition of exposure (hematocrit) and mediator (MRI neuroimaging) variables at concurrent times may have limited the accuracy of these analyses. Furthermore, mediation analyses necessitate rigorous accounting for confounders, thus observational designs may have inherent limitations in disentangling anemia’s direct causal impact on cerebral small vessel disease and dementia. Thus, future observational and translational work will be needed to assess whether anemia is an upstream risk factor that can lead to differential changes in cerebral small vessel disease burden over time, thereby impacting dementia risk.

Our study strengths included the analysis of a large, prospective population-based cohort of ethnically diverse participants, the long-term follow-up of cognitive assessments across the study course, the use of standardized MRI neuroimaging analyses, and adjustment for relevant covariates for dementia. However, several study limitations require mention. First and foremost, our study may have been subject to unmeasured confounding. While we adjusted for traditional risk factors in all models including confounders for hematocrit’s relationship with dementia (including renal function, which has previously been identified as a relevant risk factor for dementia^23,24^), we did not have data on etiologic drivers for low hematocrit/anemia (ie., inflammation, iron deficiency, nutritional status, impaired erythropoiesis). Because certain drivers for anemia are also known to relate to dementia risk, it remains unclear whether low hematocrit’s relationship with dementia was driven by hematocrit itself or underlying drivers for anemia development. Further studies assessing hematocrit as well as underlying drivers for anemia will allow for a deeper understanding of mechanistically causal relationships that can be either used as screening risk factors or even targets for dementia prevention. Second, it similarly remains uncertain whether and how low hematocrit (or its underlying drivers) directly cause dementia. Our MRI data did not appear to suggest that cerebral small vessel disease mediates this relationship. However, as previously stated, further work assessing serial neuroimaging as well as other MRI neuroimaging markers (ie., cerebral perfusion and connectivity) will be helpful in clarifying how low hematocrit impacts brain health. Additionally, human observational data will need to be paired with translational and pre-clinical models to disentangle the natural overlap of low hematocrit concentrations and underlying medical disease to establish causal relationships of low hematocrit and dementia. Lastly, our study was limited to hematocrit as the assessment of RBC concentration and RBC characteristics. It is unclear whether other factors related to the RBC number, size, maturity, or deformability of the RBC itself could play a role in dementia risk.

## CONCLUSION

We identified relationships between low hematocrit concentrations with incident dementia risk in an ethnically diverse population-based study. The biological underpinnings behind these findings are unknown. Further investigation into RBC concentrations’ impact on neuroimaging markers of brain health and dementia risk is required to assess whether this can be leveraged as a screening tool or even modifiable treatment target.

## Acknowledgement

Special thanks to the research coordinators and research participants that made this study possible.

## Author contributions

DR: conception and design of study, analysis of data, drafting manuscript

ML: analysis of data, drafting manuscript/figures

KS: acquisition of data, drafting manuscript

SA: acquisition of data, drafting manuscript

VG: acquisition of data, drafting manuscript

BL: acquisition of data, drafting manuscript

SS: conception of study, drafting manuscript

TR: acquisition of data, drafting manuscript

CW: acquisition of data, drafting manuscript

ME: acquisition of data, drafting manuscript

JG: design of study, analysis of data, drafting manuscript

## Data availability

Data supporting the findings of this study are available upon reasonable request.

Supplemental figure: Association of hematocrit with incident dementia and mediation analyses

Legend: OR: odds ratio; OR’: mediation odds ratio regressing exposure and mediator together; SVD: small vessel disease SPVS: small perivascular space; WMH: white matter hyperintensity; CMB: cerebral microbleed

